# A new targeted capture method using bacterial artificial chromosome (BAC) libraries as baits for sequencing relatively large genes

**DOI:** 10.1101/302760

**Authors:** Kae Koganebuchi, Takashi Gakuhari, Hirohiko Takeshima, Kimitoshi Sato, Kiyotaka Fujii, Toshihiro Kumabe, Satoshi Kasagi, Takehiro Sato, Atsushi Tajima, Hiroki Shibata, Motoyuki Ogawa, Hiroki Oota

## Abstract

To analyze a specific genome region using next-generation sequencing technologies, the enrichment of DNA libraries with targeted capture methods has been standardized. For enrichment of mitochondrial genome, a previous study developed an original targeted capture method that use baits constructed from long-range polymerase chain reaction (PCR) amplicons, common laboratory reagents, and equipment. In this study, a new targeted capture method is presented, that of bacterial artificial chromosome (BAC) double capture (BDC), modifying the previous method, but using BAC libraries as baits for sequencing a relatively large gene. We applied the BDC approach for the 214 kb autosomal region, *ring finger protein 213*, which is the susceptibility gene of moyamoya disease (MMD). To evaluate the reliability of BDC, cost and data quality were compared with those of a commercial kit. While the ratio of duplicate reads was higher, the cost was less than that of the commercial kit. The data quality was sufficiently the same as that of the kit. Thus, BDC can be an easy, low-cost, and useful method for analyzing individual genome region with substantial length.

## Introduction

The high-throughput sequencing technology, next-generation sequencing (NGS), has made a striking impact on genomic research and the entire biological field. The NGS technology is often called massively parallel sequencing because it effectively conducts whole-genome sequencing in a relatively short time [1]. NGS enables researchers to analyze the whole human genome of about 3 Gbp and identify all of the 30,000 genes in only 1 week [2]. To analyze specific regions (e.g., whole exons, and already known disease-related genes) using NGS, enrichment of DNA libraries with targeted capture methods are standardized.

In capture methods, probes for enriching the targeted genomic regions are called “baits” that attract molecules of interest as in fishing. There are two major approaches for relatively large-scale genomic-region enrichment, the “on-array” and “in-solution” methods. Both of these approaches target sequences up to several hundred kbp. In the on-array capture method (Roche NimbleGen products), microarrays immobilize baits that hybridize with the targeted regions and are used to enrich the genomic region of interest. Meanwhile, the in-solution method (e.g., Roche, Illumina, Agilent Technologies, and MYcroarray products) use biotinylated DNA or RNA baits to enrich targeted region. Because DNA-RNA hybrids show higher efficiency than do DNA-DNA hybrids, RNA baits are used in some systems [3]. The targeted DNA is recovered using streptavidin-labeled magnetic beads. The in-solution approach has advantages compared with the on-array method: the reagent cost is lower, less DNA is required, and it is easily scaled because the capture method can be conducted entirely in small tubes [4].

Small-scale targeted capture methods have been proposed for the enrichment of the complete mitochondrial genome (mtDNA) [5–7]. Maricic et al. (2010) presented a capture method for the mtDNA molecules that used biotinylated polymerase chain reaction (PCR) amplicons as baits. Human mtDNA is approximately 16.6 kbp long. When constructing the baits, two primer sets of long-range PCR that amplify >9 kbp regions are sufficient to cover whole mtDNA genome sequencing. The long-range PCR amplicons are sheared with sonicators. The sheared amplicons are biotinylated and used for the enrichment. This targeted capture method is cited by approximately 200 previous studies that analyzed genomes of modern or ancient organisms (e.g., humans, pathogens, animals, and fishes). The commercial targeted capture kits for small regions cost approximately 250–900 USD per reaction. The method provided by Maricic et al. (2010) is approximately 50 USD per reaction and much less expensive than commercial methods. If the Maricic’s method can be applied for large genes, then it definitely saves the cost.

In order to enrich larger genomic regions than mtDNA, we conceived of using bacterial artificial chromosome (BAC) libraries as baits, instead of PCR amplicons. BAC is a vector that can carry DNA fragments of >300 kbp [8], and can be amplified by culturing *E. coli* harboring the BAC. Human BAC libraries constructed in previous studies [9–11] are available and distributed through resource centers. We named the novel approach presenting in this study as “BAC double capture (BDC) method.” Here we show the conditions optimized for the BDC method, and the satisfactory efficiency evaluated in comparison to the commercial enrichment kit in the NGS output data.

## Materials and Methods

### Preparation of indexed libraries for testing experimental conditions by BAC single capture (BSC) and BAC double capture (BDC) with PrimeSTAR

A DNA solution purchased from the Health Science Research Resources Bank (Osaka, Japan) was used in the present study. The DNA concentration was measured using a NanoPhotometer (Implen; CA, USA). The total amount of 5 μg of DNA was sheared using a Covaris S2 sonicator (Covaris; MA, USA). The target peak was set at 400 bp. A total of 50 ng of DNA was used to produce the indexed library, using an NEBNext Ultra DNA Library Prep Kit and Multiplex Oligos for Illumina (New England BioLabs; MA, USA). Sheared DNA was end-repaired, dA-tailed, and ligated to Illumina specific adaptors. Sizes of the adaptor-ligated DNAs are selected to an approximate insert size, 400–500 bp, by Agencourt AMPure XP beads (Beckman Coulter; CA, USA). The genomic DNA shotgun library was amplified with 4 PCR reactions using a primer pair, Sol_bridge_P5 and Sol_bridge_P7, which was as presented in Maricic et al. (2010). We used 500 pmol of the library as a template for PCR in a 50-μL solution containing deoxynucleotide (dNTP) 0.2 mM, 0.2 μM of each primer, 1.25 U of PrimeSTAR GXL DNA Polymerase (Takara Bio; Shiga, Japan). PCR was carried out following the cycling reaction: 15 cycles of denaturation at 98°C for 10 sec, annealing at 60°C for 15 sec, extension at 68°C for 50 sec. Those PCR products were pooled and the solution was purified using a MinElute PCR Purification Kit (Qiagen; Hilden, Germany), and it was then eluted it into 23 μL buffer EB (Qiagen). The concentration of the solution was measured using a NanoPhotometer (Implen). The total amount of 2μg per capture reaction was obtained.

### Bait production

The BAC from the CHORI-17 library (ID number: CH17-24F19) included *RNF213* and nearby four genes with the intergenic regions (Fig 1). The total length of the BAC, from the BACPAC Resources Center (https://bacpacresources.org), was 213,477 bp. NucleoBond BAC 100 (Macherey-Nagel; Düren, Germany) was used to purify the BAC. The concentration was measured using a NanoPhotometer (Implen). The total amount of 5 μg of BAC was sheared using a Covaris S2 sonicator (Covaris). Because Maricic et al. (2010) recommended that smear DNA band of a gel electrophoresis should be brightest at a size smaller than 1 kbp, and no fragment longer than 5 kbp should be visible, four default settings of the peaks were selected: 150 bp, 300 bp, 500 bp, and 800 bp. The seven baits we obtained showed different peaks: 151 bp, 340 bp, 456 bp, 492 bp, 522 bp, 619 bp, 735 bp, and 882 bp that were included in the range of that Maricic et al. (2010) showed. The sheared BACs were purified using a MinElute PCR Purification Kit (Qiagen). Subsequently, 1.5-μg sheared BACs per capture reaction were prepared, and the products were then biotinylated according to the protocol used in the previous study [7]. To evaluate the effects of the baits’ lengths, the five baits (150 bp, 340 bp, 619 bp, 735 bp, and 882 bp) were used; for the number of the captures, two baits (340 bp or 456 bp peak) were used; and for the hybridization temperature, baits that showed a 492 bp peak were used. Baits of 522 bp peak were used for an initial BDC (with PrimeSTAR), and baits of 492 bp peak were used for an additional BDC.

**Fig 1.**
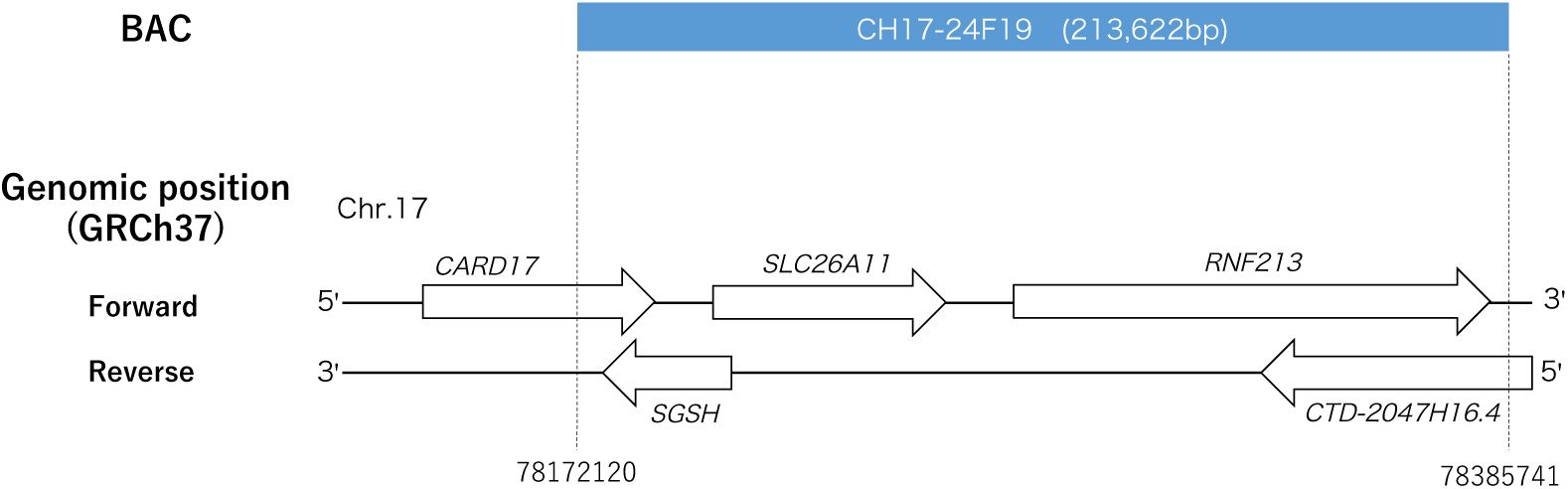
A genomic position of the BAC (ID: CH17-24F19). The BAC contains five genes: *caspase recruitment domain family member 17* (*CARD17*), *solute carrier family 26 member 11* (*SLC26A11*), *ring finger protein 213* (*RNF213*), *N*-*sulfoglucosamine sulfohydrolase* (*SGSH*), *CTD*-*2047H16.4* (uncharacterized gene).

### BAC single capture (BSC) for testing experimental conditions

This enrichment was conducted according to the protocol of Maricic et al. (2010) using BAC baits. We named it “BAC single capture (BSC).” Concentrations of enriched libraries were assessed using a KAPA Quantification Kit (Kapa Biosystems, Cape Town, South Africa). The size distributions of enriched libraries were verified using a 2100 Bioanalyzer (Agilent Technologies; CA, USA). To determine the technical variability in targeted captures, each capture was performed in duplicate.

### Initial protocol of BAC double capture (BDC) with PrimeSTAR

This enrichment was conducted following the modified protocol of the NimbleGen technical note “Double Capture: High Efficiency Sequence Capture of Small Targets for use in SeqCap EZ Library, Applications on 454 Sequencing Systems” (Fig 2). SeqCap EZ Hybridization and Wash Kit (Roche; Basel, Switzerland) were used according to the technical note. The protocol was named, “BAC double capture (BDC) with PrimeSTAR.” Blocking oligonucleotide solutions and human Cot-1 DNA were added to the library solution. The solution was dried out using a heat block at 95°C. Hybridization buffer and formamide added to the dried DNA, and the mixture was suspended by vortex mixing. The suspended mixture was single-stranded using a heat block at 95°C for 10 min. Biotinylated BAC baits (500 ng) eluted by 4.5 μL PCR grade water was added to the single-stranded DNA mixture and mixed by pipetting. Then the solution was heated in a thermal cycler to 95°C for 10 min and incubated at 65°C overnight (12–16 h). Following incubation, Dynabeads M-270 Streptavidin (Invitrogen; CA, USA) was added to the hybridization mixture. Bound DNA fragments were washed and eluted using NGS MagnaStand (Nippon Genetics, Tokyo, Japan). After the wash and the elution, PCRs were run of the enriched library (26 μL) before removing magnetic beads. The 5 μL of the eluted library was used as a template for PCR in a 50 μL solution containing deoxynucleotide (dNTP) 0.2 mM, 0.2 μM of each primer, Sol_bridge_P5 and Sol_bridge_P7 in Maricic et al. (2010), 1.25 U of PrimeSTAR GXL DNA Polymerase (Takara Bio). The 1st post-capture PCR was carried out following the cycling reaction: 16 cycles of denaturation at 98°C for 10 sec, annealing at 60°C for 15 sec, extension at 68°C for 50 sec into the plateau phase according to Maricic et al. (2010). The PCR amplicon was purified using a MinElute PCR Purification Kit (Qiagen). Then, the 2nd capture was conducted using the enriched and purified library using the same steps as in the 1st capture. After that, the 2nd post-capture PCR of the 2nd captured library was run into the plateau phase (20 cycles) using the same cycling condition as in the 1st post-capture PCR. The amplified library was purified using the methods described above. Quantification of the amplified capture library was conducted with a KAPA Library Quantification Kit for Illumina NGS platforms (Kapa Biosystems) and a 2100 Bioanalyzer (Agilent Technologies). To determine the technical variability in targeted captures, each capture was performed in duplicate.

**Fig 2.**
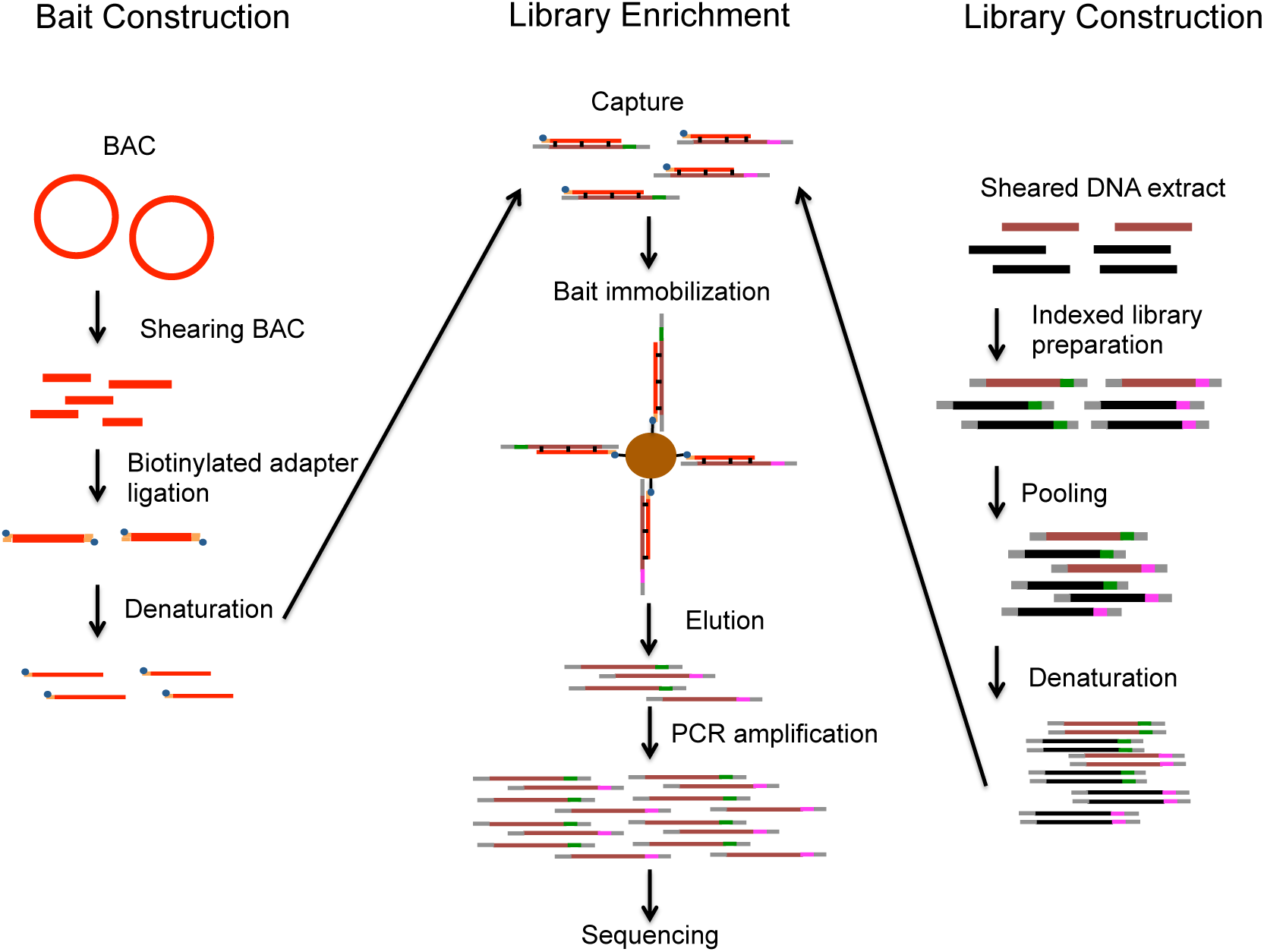
An overview of the BAC double capture (BDC) method, which we modified (Maricic et al., 2010). On the left, the bait construction from the BAC is shown; on the right, the production of indexed libraries that are used in the library enrichment (center). Those colored light red are the BAC-based baits, dark red represents targeted DNA molecules in the libraries, black represents non-targeted DNA molecules in the libraries, green and pink represent indexes, gray represents adapters, and blue and yellow represent biotinylated adapters. Thick lines represent double stranded DNA, and thin lines represent single stranded DNA.

### Sequencing for BSC and BDC with PrimeSTAR

The enriched libraries by BSC were sequenced on a MiSeq (Illumina; CA, USA) using Illumina MiSeq reagent kit v2 (2 × 25 cycles) or v2 nano (2 × 150 cycles) or v3 (2 × 75 cycles). Fastq files were processed using Trimmomatic (version 0.35) in the paired-end palindrome mode to remove TruSeq adapter sequences, low-quality reads (average: <Q20), and nucleotides after the 5’-end from the 26th base and following bases of each read, regardless of quality, to minimize the differences among the three reagent kits.

The enriched libraries by BDC with PrimeSTAR were sequenced on a MiSeq (Illumina) using the Illumina MiSeq Reagent Kit v3 (2 × 75 cycles). Fastq files were processed using Trimmomatic (version 0.35) in the paired-end palindrome mode to remove TruSeq adapter sequences and low-quality (average: <Q20) reads.

### Alignment for BSC and BDC with PrimeSTAR

The quality-controlled reads were aligned with the Burrows-Wheeler Aligner (BWA) software version 0.7.12-r1039 [12] to the human genome (GRCh37) with default parameters. Duplicate reads per sample were marked using the MarkDuplicates tool from the Picard software version 1.128 (https://broadinstitute.github.io/picard/) and local realignments around indels were performed on per sample basis using the IndelRealigner tool from the Genome Analysis Toolkit (GATK) software version 3.4-46 [13]. The reads mapped to a large tandem repeat, chr17: 78234665-78372586, were removed. Coverage and average depth per sample of targeted regions were calculated using GATK’s DepthOfCoverage analysis. The number of mapped and duplicated reads were obtained using SAMtools version1.2 flagstat analysis [14].

### Production of indexed libraries for BDC and MB

DNA was extracted from bloods of 24 moyamoya disease (MMD) cases collected at Kitasato University Hospital using DNA Extractor WB Kit (Wako Pure Chemical Industries; Osaka, Japan). All the patients included in this study provided written informed consent. This project was approved by the ethics committee at Kitasato University School of Medicine. The concentrations of DNA extracts were measured using a NanoPhotometer (Implen) and Qubit 3.0 Fluorometer (ThermoFisher Scientific; MA, USA). Using a Covaris S220 sonicator (Covaris), 2 μg of DNA was sheared. The target peak was set at 300 bp. To produce an indexed library using NEBNext Ultra DNA Library Prep Kit and Multiplex Oligos for Illumina (New England BioLabs), 500 ng of DNA was used. Sheared DNA was end-repaired, dA-tailed, ligated to Illumina specific adaptors, size selected to an approximate insert size of 400–500 bp by Agencourt AMPure XP beads (Beckman Coulter), and amplified by 6 or 7 cycles of PCR. The libraries were purified using Agencourt AMPure XP beads (Beckman Coulter).

### The final protocol of BDC

The protocol of BDC with PrimeSTAR was modified with KAPA HiFi DNA Polymerase. The following is the final protocol of BDC. The PCR amplification process of the protocol of BDC was improved with KAPA HiFi DNA Polymerase. DNA libraries of eight MMD cases were used for the final protocol. The 1st and 2nd post-captured libraries were used as templates for PCR in a 50 μL solution containing deoxynucleotide (dNTP) 0.3 mM, 0.5 μM of each primer, 1.0 U of KAPA HiFi DNA Polymerase (Kapa Biosystems). PCR was carried out using the following protocol: an initial denaturing step at 98°C for 2 min, 8 cycles for 1st post-capture libraries or 13 cycles for the 2nd post-capture libraries of denaturation at 98°C for 20 s, annealing at 60°C for 30 s, extension at 72°C for 40 s, and a final extension step at 72°C for 5 min. To determine the technical variability in targeted captures, each capture was performed in duplicate.

### MYbaits double capture (MB)

DNA libraries of 24 MMD cases were used for the targeted capture experiment. The captures were performed using the MYbait Custom Kit (MYcroarray; MI, USA) constructed for enrichment of the same region of the BAC twice, following the manufacturer’s instructions (http://www.mycroarray.com/pdf/MYbaits-manual-v3.pdf). The baits of MYbaits were uniquely designed to map to the human reference genome. The designed baits covered nearly 80% of the target region. The libraries were hybridized to half of an aliquot of the RNA baits per reaction. The 1st post-capture PCR was 8 cycles. The 2nd capture was conducted using the whole quantity of the 1st post-capture PCR amplicons with the same protocol as that in the 1st capture. The 2nd post-capture PCR was 11 cycles. After the enrichment, the libraries were purified using MinElute PCR Purification kit (Qiagen), quantified using TapeStation (Agilent Technologies) and Qubit 3.0 Fluorometer (ThermoFisher Scientific). The method is called MYbaits double capture (MB).

### Sequencing and alignment for BDC and MB

The enriched libraries were sequenced on a MiSeq (Illumina) using Illumina MiSeq reagent kit v3 (2 × 75 bp chemistry). Fastq files were processed using Trimmomatic (version 0.35) [15] in the paired-end palindrome mode to remove TruSeq adapter sequences and low-quality (average: <Q20) reads. The method of alignment for BDC and MB was the same as for that of BSC and BDC with PrimeSTAR.

### Valiant calling for BDC and MB

The resulting data was analyzed with the GATK version 3.4-46, according to GATK Best Practices recommendations [13,16,17]. Following the guidelines for experiments of small-targeted regions, this workflow included calling variants and producing the genomic variant call format (gVCF) files in target regions individually per subject using a HaplotypeCaller, followed by joint genotyping data to produce a multisample raw VCF file using GenotypeGVCFs. Default settings were used for both tools. After variant calling, the following annotations and thresholds were used to remove low-confidence SNPs, based on GATK recommendations for hard filtering: QD <2.0, FS >60.0, HaplotypeScore >13.0, MQ <40, MQRankSum <–12.5, ReadPosRankSum <–8.0. Similarly, the following filters were applied to remove low-confidence indels: QD <2.0, FS >200.0, ReadPosRankSum < – 20.0. We extracted variants information from the filtered VCF file using VCFtools [18].

## Results

### Experimental conditions of BDC

The effects of baits lengths were evaluated. BSC was performed with the five baits with different lengths and the on-target rates were graphed (Fig 3). The approximate 350–750 bp peak baits showed more stability and higher on-target rates (0.28–0.43%) than did the 150 bp peak bait (0.16% and 0.33%). The rates of the 150 bp peak bait showed a larger difference (0.17%) between the duplicates than did that of the longer baits (0.04–0.08%).

**Fig 3.**
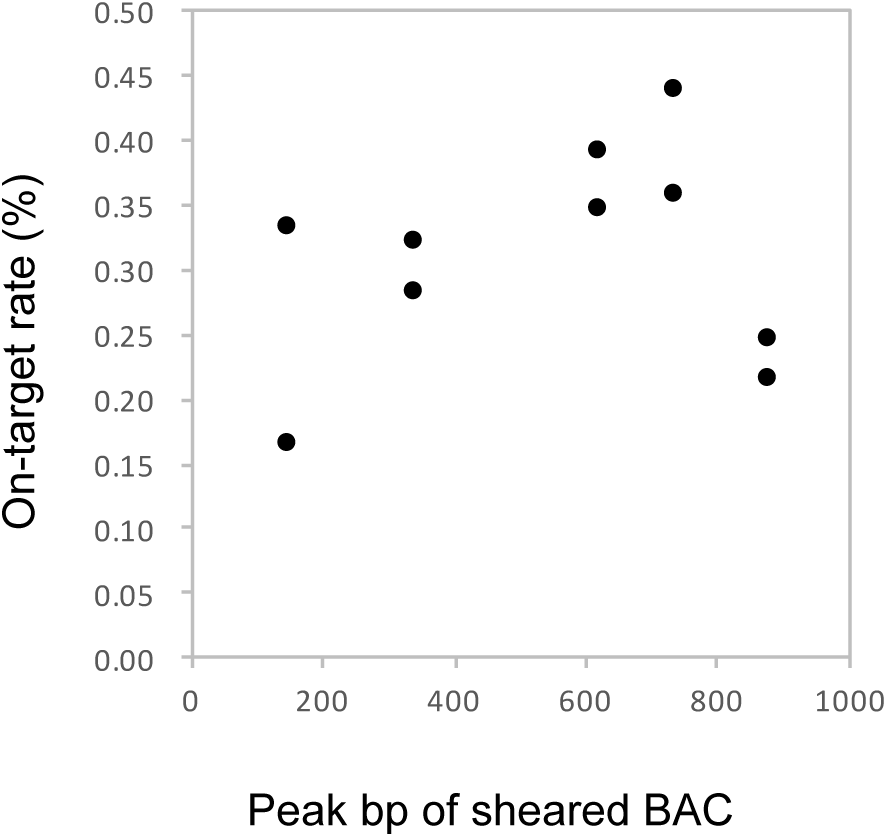
On-target rates depending on BAC baits length. On-target rate equals the reads mapped to the target region divided by the reads mapped to the whole reference genome.

The effects of numbers of captures were evaluated. BSC and BDC were performed with PrimeSTAR and the on-target rates were graphed (Fig 4). There was one capture of BSC and two of BDC with PrimeSTAR, which showed higher on-target rates (16.53% and 12.21%) than did that of BSC (0.28% and 0.32%). Therefore, two captures were more efficient than one.

**Fig 4.**
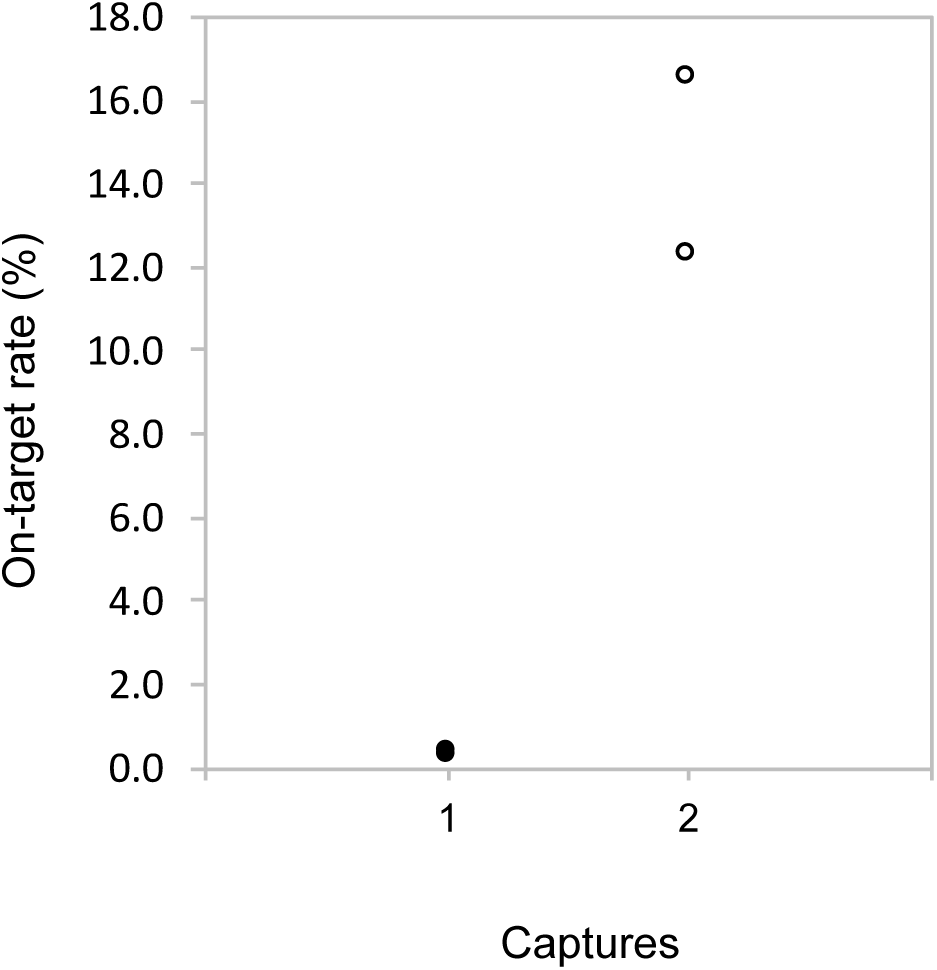
On-target rates depending on the number of captures. The formula to calculate the on-target rate is the same as in Fig 3.

The effects of hybridization temperature were evaluated. Hybridizations were performed at 45°C and 65°C, and the on-target rates were graphed (Fig 5). Hybridization at 65°C showed higher on-target rates (21.12% and 25.01%) than did those hybridized at 45°C (3.51% and 5.78%).

**Fig 5.**
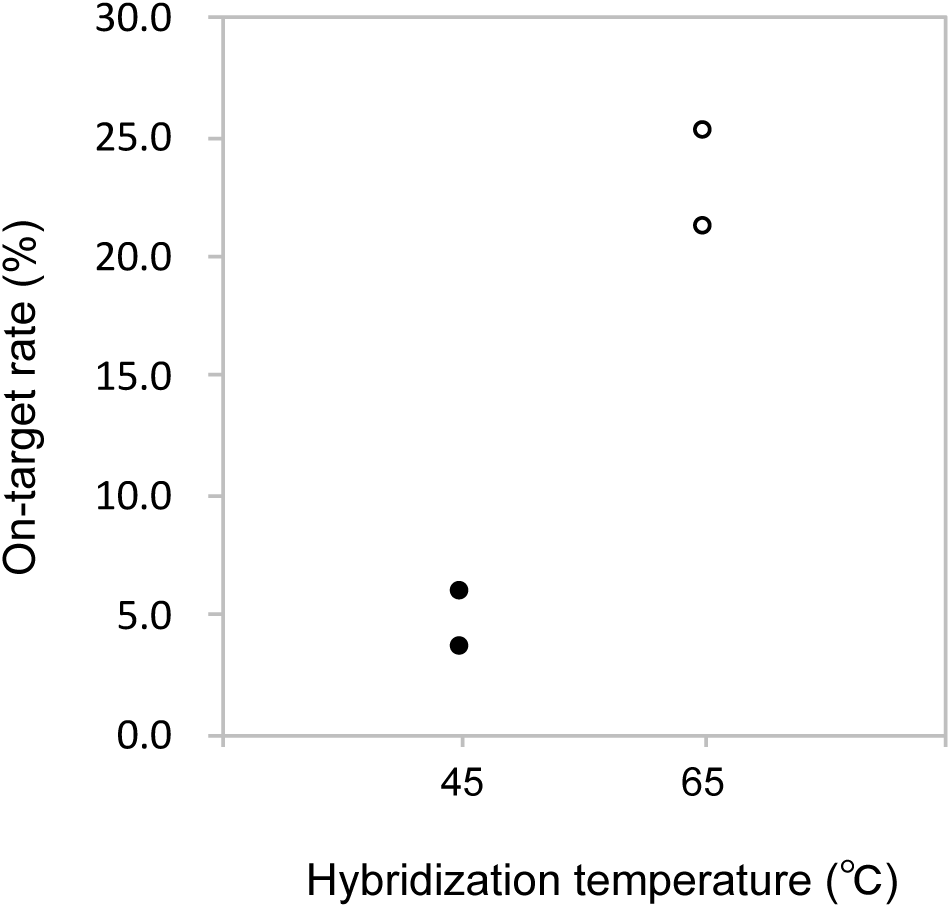
On-target rates of the hybridization temperature. The formula to calculate the on-target rate is the same as in Fig 3.

### Comparison between BDC and a commercial targeted capture method

PrimeSTAR GXL DNA polymerase for BDC was used in the initial protocol of BDC (see Materials and Methods). However, in the final protocol of BDC, it was converted to KAPA HiFi DNA polymerase (see Materials and Methods) because the polymerase showed more high yield than did the PrimeSTAR GXL DNA polymerase (S1 Figure, S1 Protocol). The quality of the NGS data of BDC with KAPA HiFi DNA polymerase and the targeted capture method was compared with that of MB.

Averages of rates of “unique reads” and “duplicate reads” were calculated (Table 1). “Unique reads” mean reads that mapped uniquely to a reference genome. “Duplicate reads” mean reads that mapped to a reference genome at the same position with the other reads and had the same length and the same variation. When raw NGS datasets are processed, duplicate reads are removed from the dataset. BDC with KAPA HiFi DNA polymerase showed a higher average of duplicate-reads rate (46.9%) than did that of MB (16.1%). The averages of depths between BDC and MB were compared (Table 1). “Depth” means the number of reads that mapped at a genomic position. The averages of total depth calculated by adding depth of each genomic position were almost the same, approximately 30 M. The averages of depth were also almost the same, approximately 140. The averages of on-target rates were also compared (Table 1): that of BDC (22.5%) was similar to that of MB (24.3%).

**Table 1.** Average numbers of BDC and MB.

| **Method** | **BDC** | **MB** |
| --- | --- | --- |
| **Duplicate reads (%)** | 46.9 | 16.1 |
| **Unique reads (%)** | 53.1 | 83.9 |
| **Total depth** | 30,020,317 | 30,321,836 |
| **Depth** | 140.5 | 141.9 |
| **On-target rate (%)** | 22.5 | 24.3 |

### Validation of variant sites

The validation of BDC was evaluated using sequence data of 8 samples from MMD cases that were conducted both BDC and MB. The called SNP sites of BDC (572 sites) were larger than those of MB (549 sites) (Table 2). The numbers of SNPs registered in dbSNP were larger in BDC (540 sites) than in MB (517 sites). The number of SNPs not registered in dbSNP of BDC was the same as that of MB. The concordant rates of genotypes of SNP sites between BDC and MB were calculated (Table 3). The SNPs registered in dbSNP were 98.4%. The SNPs registered in dbSNP were 97.3%. In twenty-eight SNPs, at least 1 out of 8 MMD samples were called different genotypes between BDC and MB. Those SNPs were placed at genomic regions where was difficult to map reads and call variants correctly in (poly A or G regions: 7 sites, CNV: 11 sites, retro transposons: 8 sites, low complexity regions: 2 sites).

**Table 2.** Numbers of validated SNPs.

| **Method** | **All SNPs** | **dbSNP (registered)** | **dbSNP (non-registered)** |
| --- | --- | --- | --- |
| **BDC** | 572 | 540 | 32 |
| **MB** | 549 | 517 | 32 |

**Table 3.** Concordant rates of genotypes.

|  | **dbSNP (registered)** | **dbSNP (non-registered)** |
| --- | --- | --- |
| **Number of sites\*** | 478 | 32 |
| **Concordant rate of**<br>**Genotypes** | 98.4% | 97.3% |

\*Sites that genotyped all eight samples with both BDC and MB.

### Comparisons of the average values of between BDC and MB

The data qualities and costs of BDC and MB were evaluated comparing nine categories (Table 4). The required genomic DNA for each method was higher weight in BDC (1.5 μg) than that in MB (0.5 μg). BDC required baits which we constructed myself, while MB required manufactured baits included in a targeted capture kit. BDC took 10 days to prepare the baits because of the time required to order *E. coli* harboring BAC that covered *RNF213* and amplified and purified it. MB took 60 days from design of the baits to its arrival. The period of a targeted capture experiment in BDC was the same as MB, 3 days. The experimental cost without sequencing using MiSeq of BDC (USD 55) was lower than that of MB (USD 270). We examined four points regarding the quality of the data. That of MB was the required library weight. The duplicate read rate of BDC (46.9%) was higher than that of MB (16.1%). The average of depth of BDC (140.5) was the almost same as that of MB (141.9). The on-target rate of BDC (22.5%) was also quite close to that of MB (24.3%). Therefore, the data quality of BDC was close in on MB, except only for the duplicate read rate.

**Table 4.** Summary of comparisons between BDC and MB.

| Category |  | Method |  |
| --- | --- | --- | --- |
| 1 | 2 | BDC | MB |
| Cost | Required library weight (µg) | 1.5 | 0.5 |
|  | Baits construction | Self-making | Outsource |
|  | Preparation period of baits (day) | 10 | 60 |
|  | Period of targeted capture (day) | 3 | 3 |
|  | Cost per reaction (USD) | 55 | 270 |
| Data<br>quality | Duplicate reads (%) | 46.9 | 16.1 |
|  | Average of depth | 140.5 | 141.9 |
|  | On-target rate (%) | 22.5 | 24.3 |
|  | SNP concordance* (%) | 98.4 |  |

\*A calculated value from SNPs registered in dbSNP138

## Discussion

We compared the three experimental conditions, the baits lengths, the number of captures, and the hybridization temperatures, and found appropriate conditions exclusively for the BDC method. These might not be the best conditions, but better ones for the method.

We first found that the baits of 350–750 bp peak obtained a higher on-target rate than did the peak baits around 150 bp and 900 bp (Fig 3). The magnetic beads (Dynabeads M-270) that immobilize baits reduce the binding capacity for large DNA fragments due to the likelihood of steric hindrance. Twice as many copies of a 500 bp DNA fragment bind to the beads than in the case of a 1,000 bp DNA fragment. Therefore, around 900 bp baits would show lower on-target rates. The present study showed that about 150 bp baits were the lowest and most unstable on-target rates. Commercial targeted capture kits have uniformly about 100 bp baits (e.g., Agilent: 120 bp, Illumina: 100 bp, MYcroarray: 80–120 bp). The protocols of such kits optimized many conditions (e.g., bait lengths, the bait densities, the hybridization temperatures, and the hybridization reagents). Those results indicate that the experimental conditions of the protocol in the present study are not applicable for the short baits. To optimize the baits lengths for BDC, we should examine more patterns of baits lengths in the next step.

We observed much higher on-target rates with double captures than that with a single capture (Fig 4), suggesting that the double capture definitely enriches more target libraries. The targeted captures of small regions are especially more difficult than those of larger regions, because genome libraries have smaller volumes of narrow targeted regions (e.g., hundreds kbp) than those of broad targeted regions (e.g., several Mbp). Double captures seem to enable researchers to enrich more DNA of interest than do single captures.

We found that the hybridization temperature at 65°C was suitable for effective library enrichment (Fig 5), suggesting that the hybridization at 65°C gave higher specificity than that at 45°C. From these results, overall, we would propose that the conditions are the baits with 350–750 bp peak, twice capturing, and the 65°C hybridization temperature.

The average of duplicate read rate of BDC was higher than that of MB (Table 1). In the BDC protocol, PCR includes one more cycle than that in the MB protocol for obtaining libraries of adequate quantities. This could be the reason why the higher rate of duplication in BDC was observed. It might be important for improving BDC to reduce the number of PCR cycles.

The on-target rate of BDC (Table 1) was higher than the that of the previous manual methods that enriched mtDNA using baits constructed from long-range PCR amplicons or genomic regions using BAC-based baits [7,19]. A previous study [7] proposed a single capture method. Another previous study [19] also proposed a single capture method and used non-sheared BACs that were affected by their own steric hindrance. Thus, we would claim that BDC is improved from the previous manual capture methods.

BDC showed more called SNP sites than MB (Table 2). The baits used in MB were synthetic oligonucleotide probes. Non-unique baits in a human-reference-genome sequence were excluded in the probes designed. Therefore, the baits of MB covered approximately 80% of the target region. On the other hand, the baits used in BDC were constructed from BAC, which covered the whole target regions. The differences of processes constructing probes between BDC and MB could affect the numbers of enriched libraries and the called SNP sites.

The concordance of SNP genotypes between BDC and MB was >97% (Table 3). The SNPs that were called the different genotypes placed at poly A, poly G, repetitive regions and transposons. Those genomic regions that showed low uniqueness in genomes are, in general, more likely to have errors in PCRs and mapping reads [20]. Thus, the SNP sites that showed discordance of genotypes locate genomic regions where were difficult to call SNPs correctly.

BDC allows efficient capture of the genomic library of NGS for large genes. First, the approach is cost-effective in that it only requires standard laboratory equipment and reagents that cost USD55 per reaction (Table 4). Second, it is fast, i.e., it enables researchers to perform captures immediately without designing and constructing baits (Table 4). Third, the on-target rates are almost the same for BDC and MB (Tables 1 and 4). Those features enable more laboratories to start easily targeted captures. More adjustments of capture conditions make a better BDC. When BDC is used for the other human genomic regions and any organism’s genome, the conditions of targeted captures may require adjustment. E.g., if the GC (guanine and cytosine) contents of targeted regions are different from those of the BAC used in the present study, suitable hybridization temperatures can be changed. BDC enables the recovery of targeted genomic regions like a large gene from most such ancient samples. Because a large amount of non-targeted DNA and bacterial DNA extracted from those bones, those samples are needed to retrieve only endogenous genomes. Our new approach helps to conduct paleogenomic studies.

## Acknowledgments

We thank the members of the Laboratory of Evolutionary Anthropology, especially Dr. Shoji Kawamura and Dr. Minoru Yoneda (The University of Tokyo), for many discussions and suggestions about our research. Alumni of the Department of Anatomy, Dr. Takafumi Katsumura (Okayama University) and Dr. Hiromi Matsumae (Yokohama City University), have given us great suggestions for experiments and helped with data analyses. Dr. Akiyoshi Takahashi (Kitasato University) helped us with our targeted capture experiments. Dr. Masahira Hattori, Dr. Kenshiro Oshima (The University of Tokyo) gave us equipment making a genome library for next-generation sequencing. Dr. Alison Devault at MYcroarray provided a lot of advice to construct the bait set. And we thank Robert E. Brandt, Founder, CEO, and CME, of MedEd Japan, for editing and formatting the manuscript.

## Supporting information

### Supplementary Figure Legends

**S1 Fig.**
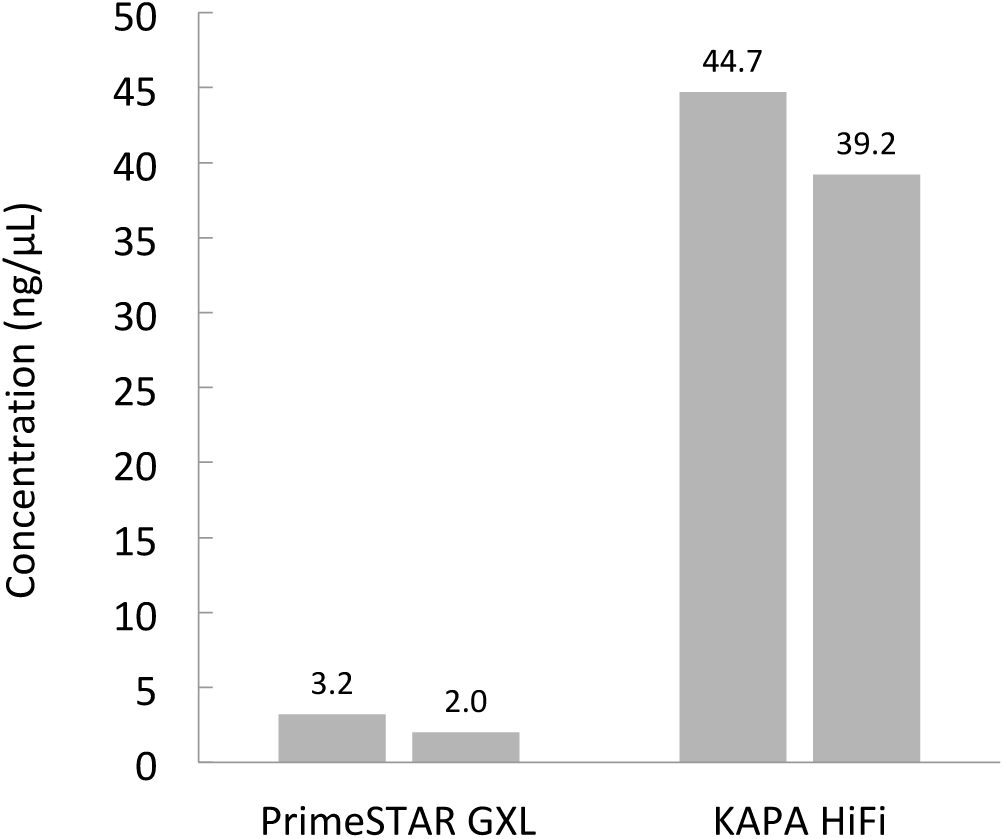
Comparison of PCR efficiency between PrimeSTAR GXL DNA Polymerase and KAPA HiFi DNA Polymerase.

### S1 Protocol Amplification efficiency of two polymerases

A 2 μL of 1st post-captured library solution before removing magnetic beads concentrated using a BDC method was used as a template for PCR in a 20 μL solution containing 0.5 U of PrimeSTAR GXL DNA Polymerase (Takara Bio), deoxynucleotide (dNTP) 0.2 mM, 0.2 μM of each primer, Sol_bridge_P5 and Sol_bridge_P7 in Maricic et al. (2010). The PCR and purification were carried out using the same method as BDC with PrimeSTAR. The same volume of the 1st post-captured library solution was used as a template for PCR in a 20 μL solution containing, 0.4 U of KAPA HiFi DNA Polymerase (Kapa Biosystems), deoxynucleotide (dNTP) 0.3 mM, 0.5 μM of each primer, Sol_bridge_P5 and Sol_bridge_P7 in Maricic et al. (2010). PCR was carried out using the following protocol: an initial denaturing step at 98°C for 2 min, 16 cycles for the 1st post-capture library of denaturation at 98°C for 20 s, annealing at 60°C for 30 s, extension at 72°C for 45 s, and a final extension step at 72°C for 5 min. To determine the technical variability in targeted captures, each PCR was performed in duplicate. The PCR amplicons were quantified using Qubit 3.0 Fluorometer (ThermoFisher Scientific).

